# Tumor microtubes connect pancreatic cancer cells in an Arp2/3 complex-dependent manner

**DOI:** 10.1101/841841

**Authors:** Casey J. Latario, Lori W. Schoenfeld, Charles L. Howarth, Laura E. Pickrell, Fatema Begum, Dawn A. Fischer, Olivera Grbovic-Huezo, Steven D. Leach, Yolanda Sanchez, Kerrington D. Smith, Henry N. Higgs

## Abstract

Actin-based tubular connections between cells have been observed in many cell types. Termed “tunneling nanotubes (TNTs)”, “membrane nanotubes”, “tumor microtubes (TMTs)”, or “cytonemes”, these protrusions interconnect cells in dynamic networks. Structural features in these protrusions vary between cellular systems, including tubule diameter and presence of microtubules. We find tubular protrusions, which we classify as TMTs, in a pancreatic cancer cell line, DHPC-018. TMTs are present in DHPC-018-derived tumors in mice, as well as in a mouse model of pancreatic cancer and a sub-set of primary human tumors. DHPC-018 TMTs have heterogeneous diameter (0.39 – 5.85 μm, median 1.92 μm) and contain actin filaments, microtubules, and cytokeratin 19-based intermediate filaments. The actin filaments are cortical within the protrusion, as opposed to TNTs, in which filaments run down the center of the tube. TMTs are dynamic in length, but are long-lived (median > 60 min). Inhibition of actin polymerization, but not microtubules, results in TMT loss. A second class of tubular protrusion, which we term cell-substrate protrusion (CSP), has similar width range and cytoskeletal features but make contact with the substratum as opposed to another cell. Similar to previous work on TNTs, we find two assembly mechanisms for TMTs, which we term “pull-away” and “search-and-capture”. Inhibition of the Arp2/3 complex, an actin assembly factor, inhibits TMT assembly by both mechanisms. This work demonstrates that the actin architecture of TMTs is fundamentally different from that of TNTs, as well as demonstrating the role of Arp2/3 complex on TMT assembly.

## Introduction

Cells possess a variety of mechanisms for exchange of materials and information, including soluble growth factors/chemokines, exosomes, adherens junctions, and gap junctions (Ribeiro-Rodrigues *et al.*, 2017; Mathieu *et al.*, 2019). Over 15 years ago, it was revealed that tubular connections can exist between cells. Originally termed cytonemes (Ramírez-Weber and Kornberg, 1999) or tunneling nanotubes (TNTs) (Önfelt and Davis, 2004; Rustom *et al.*, 2004), these protrusions are thin (<500 nm width) and can extend over 100 μm. Given their dimensions, and the fact that they are not on the basal surface but are extended between cells like a tightrope, the protrusions are easily destroyed by certain fixation procedures (Rustom *et al.*, 2004; Sartori-Rupp *et al.*, 2019), perhaps contributing to their relatively late identification.

Since these initial discoveries, a variety of similar structures have been described in multiple systems. With this wider spread has come a wider variety of structural features. One difference between the structures concerns cytoskeletal composition. While the initially identified structures were shown to contain actin but not microtubules (Ramírez-Weber and Kornberg, 1999; Rustom *et al.*, 2004; Sowinski *et al.*, 2008; Sartori-Rupp *et al.*, 2019), a number of subsequently identified structures contain microtubules (Önfelt *et al.*, 2006; Gerdes *et al.*, 2013; Osswald *et al.*, 2015; Kumar *et al.*, 2017; Resnik *et al.*, 2018; Kim *et al.*, 2019). Where examined, intermediate filaments have also been identified (Iglič *et al.*, 2007; Ady *et al.*, 2014; Sáenz-de-Santa-María *et al.*, 2017; Resnik *et al.*, 2018). A second difference concerns the size of the structures. TNTs are generally defined as having a diameter less than 500 nm (Rustom *et al.*, 2004; Ariazi *et al.*, 2017; Sartori-Rupp *et al.*, 2019), but other structures can be several microns wide (Vidulescu *et al.*, 2004; Osswald *et al.*, 2015; Kumar *et al.*, 2017). These variations have led to use of additional names such as ‘microtubes’ to reflect the structural differences. To avoid a diameter specification, we will refer to the structures collectively as intercellular membrane tubules (IMTs).

IMTs have been identified *in vivo* in both *Drosophila* (Ramírez-Weber and Kornberg, 1999; Roy *et al.*, 2011, 2014; Huang *et al.*, 2019) and mammals (Chinnery *et al.*, 2008) suggesting physiological significance. A number of communication functions have been attributed to IMTs, including targeted growth factor transmission (Roy *et al.*, 2011, 2014), neurotransmitter signaling from epithelial cells (Huang *et al.*, 2019), and direct exchange of cytoplasmic materials including: mitochondria (Kumar *et al.*, 2017; Kretschmer *et al.*, 2019), endosomes/lysosomes (Rustom *et al.*, 2004; Kumar *et al.*, 2017; Sartori-Rupp *et al.*, 2019), plasma membrane proteins (Önfelt and Davis, 2004), mRNA (Haimovich *et al.*, 2017), and microRNAs (Thayanithy *et al.*, 2014).

Three pathogenic roles have also been linked to IMTs. First, pathogens such as viruses (Sowinski *et al.*, 2008; Hashimoto *et al.*, 2016; Kumar *et al.*, 2017; Panasiuk *et al.*, 2018) and bacteria (Önfelt *et al.*, 2006; Kim *et al.*, 2019) can use IMTs as a means of cell-to-cell transfer. Second, mis-folded protein aggregates including huntingtin (Costanzo *et al.*, 2013), prion protein (Zhu *et al.*, 2015), and α-synuclein (Abounit *et al.*, 2016) have also been shown to transfer between cells through IMTs. Finally, IMTs have been linked to increased treatment resistance of several cancers. In a glioblastoma model, IMTs of > 100 μm extended at the invasive edge of the tumor into peripheral tissue. These IMTs had a diameter ∼1.6 μm, contained both actin and microtubules, and correlated with increased resistance to radiation and chemotherapy (Osswald *et al.*, 2015; Weil *et al.*, 2017). Similarly, IMTs that contribute to treatment resistance have been identified in pancreatic cancer (Desir *et al.*, 2018), prostate cancer (Kretschmer *et al.*, 2019), mesothelioma (Lou *et al.*, 2012) and leukemias (Polak *et al.*, 2015).

In terms of IMT assembly, there is strong consensus that the process is actin-dependent (Ariazi *et al.*, 2017). Interestingly, two distinct mechanisms for assembly have been observed (Sowinski *et al.*, 2008; Veranič *et al.*, 2008; Gerdes *et al.*, 2013; Kumar *et al.*, 2017), which we term ‘pull-away’ and ‘search-and-capture’. In pull-away assembly, an IMT is created between two cells that are closely associated when one of the cells migrates away. In search-and-capture, one cell extends a protrusion that makes stable contact with another cell. The factors that nucleate the actin filaments involved in IMT assembly by either mechanism have not been defined. Due to their thin, tubular nature and their actin dependence, it has been tempting to view IMTs as specialized filopodia. However, evidence from a neuronal cell line suggests distinct and even opposite molecular characteristics between IMTs and filopodia (Delage *et al.*, 2016), although IMTs in this system still contain parallel actin filaments that run the length of the IMT, similar to filopodia (Sartori-Rupp *et al.*, 2019).

In this study, we examine IMTs of a variety of diameters emanating from a low-passage pancreatic ductal adenocarcinoma (PDAC) cell line, which we define as TMTs based on their presence in tumors and morphological characteristics. Actin filaments are present in all IMTs, with microtubules and intermediate filaments also present in >90% of the structures. In IMTs of median width (approximately 2 μm), actin filaments are enriched along the IMT membrane whereas microtubules and intermediate filaments run down the center. A second structure, which we refer to as a cell surface protrusion (CSP), does not contact another cell but interact with the substratum. IMT assembly occurs equally by pull-away and search-and-capture mechanisms. Inhibition of Arp2/3 complex results in a decrease in IMT assembly by both mechanisms. We also find IMTs in PDAC tumors in several contexts, including a sub-set of primary human tumors. These results suggest that IMTs can represent a heterogeneous population of structures, even within a single cell type.

## Results

### DHPC-018 cells possess two types of lateral finger-like protrusions

The DHPC-018 (Dartmouth-Hitchcock Pancreatic Cancer) cell line was established from a human peritoneal metastasis that had been surgically removed after neo-adjuvant treatment. Examination of live DHPC-018 cells by differential interference contrast (DIC) microscopy revealed two clear features. First, the cells contain frequent intercellular connections, in the form of thin protrusions (Figure 1A). The mean frequency of these protrusions is 0.315±0.06 protrusions/cell (Figure 1B). A wide variety of protrusion lengths is present, with a range from 5 – 244 μm, and a median of 26.7 μm (Figure 1C). The width of the protrusions is also variable, with a range from 0.39 – 5.85 µm and a median of 1.92 μm (Figure 1D). Due to their width (generally larger than traditionally defined TNTs) and their presence in a tumor-derived cell line, we choose to refer to these structures as tumor microtubes (TMTs).

**Fig. 1.**
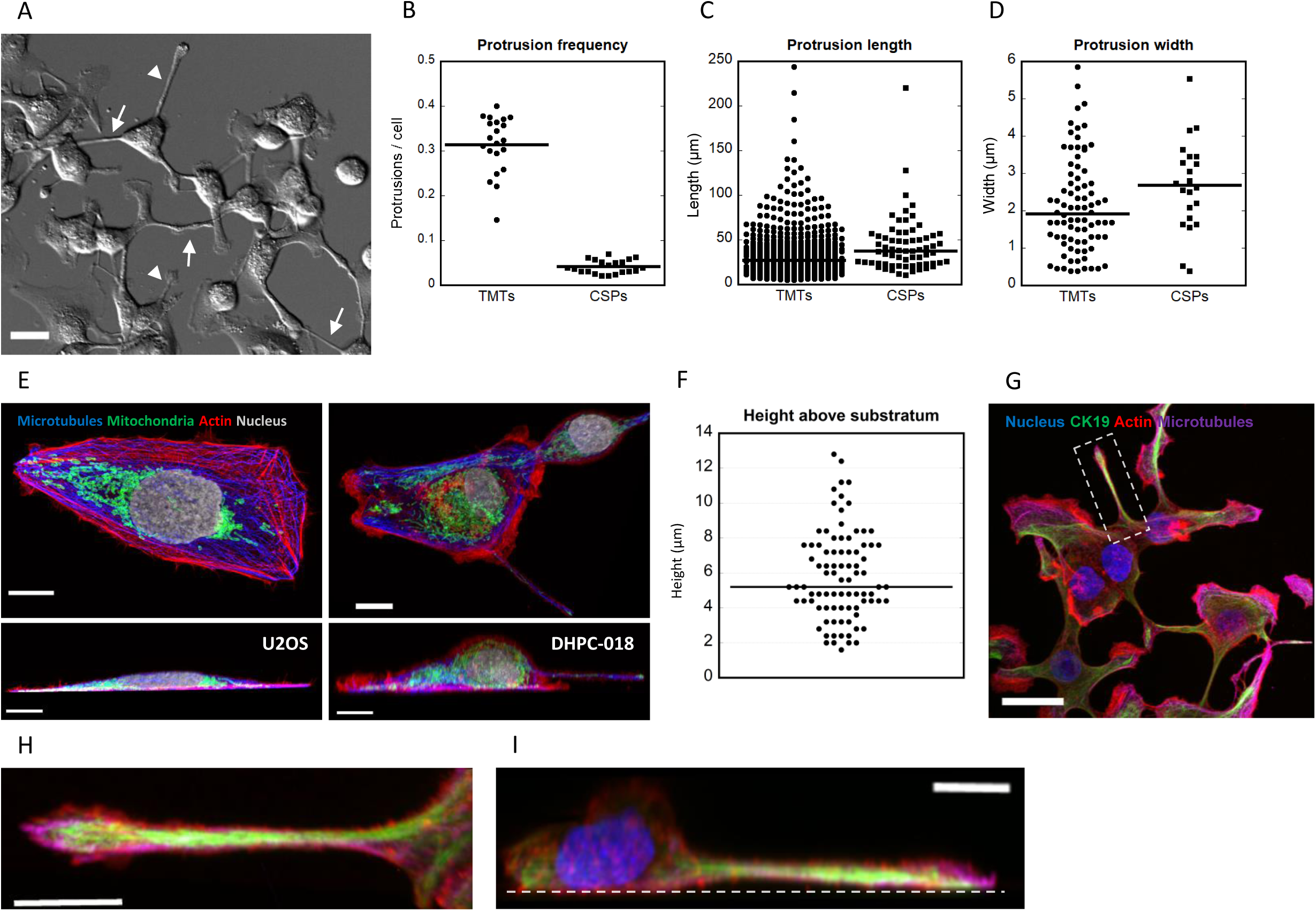
DHPC-018 cells possess intercellular membrane tubules (TMTs) and cell-substratum protrusions (CSPs) of variable length and width. (A) DIC microscopy of live DHPC-018 cells. Focal plane starts 1 µm above basal surface, 20x, 0.75 NA objective. Arrows and arrowheads denote examples of TMTs and CSPs respectively. Scale bar, 25 μm. (B) TMT and CSP frequency. 21 fields, 4963 cells, 1487 TMTs, 182 CSPs. Bars are medians: 0.315 ± 0.01 TMT/cell (SEM), 0.041 ± 0.01 CSP/cell. (C) TMT and CSP length. 544 TMTs, 63 CSPs. Bars are medians: 26.7 ± 1.2 μm for TMT (SEM), 37.7 ± 3.8 μm for CSPs. (D) TMT and CSP width. 82 TMTs, 22 CSPs. Bars are medians: 1.90 ± 0.14 μm for TMT (SEM), 2.68 ± 0.25 μm for CSPs. (E) Comparative Airyscan images of fixed U2OS (left) and DHPC-018 (right) cells, stained with anti-Tom20 (green, mitochondria), anti-tubulin (blue), TRITC-phalloidin (red), and DAPI (white). 0.2 µm Z slices, 100x 1.4 NA objective. Top: max intensity plan view. Bottom: side view of Z stack. Scale bars, 10 µm. (F) TMT height above the substratum. N = 82 TMTs. Bar is median, 5.2 ± 0.27 μm (SEM). (G) Example of CSP. Max intensity plan view of a field of DHPC-018 cells (Airyscan, 0.2 µm Z slices). Cells stained with anti-CK19 (green), anti-tubulin (magenta), TRITC-phalloidin (red), and DAPI (blue). Scale bar, 25 µm. (H) Zoom of the boxed CSP from (G). Scale bar, 10 µm. (I) Side view of Z stack of boxed CSP from (G). Dashed line indicates the substratum. Scale bar, 10 µm.

A second interesting feature is that most DHPC-018 cells do not spread extensively on the fibronectin-coated substratum, but rather remain extended in the Z direction. This feature is best appreciated in comparison to an established culture cell line such as U2OS osteosarcoma cells (Figure 1E, Supplementary Movie 1). Examination of Z-stacks of DHPC-018 cells shows that TMTs typically protrude from the lateral cell surface, at a median height of 5.2±2.5 μm above the base of the cell (Figure 1F).

We also observed a second population of protrusions from DHPC-018 cells, that did not connect with another cell but to the substratum (Figure 1A, G, H), and which we term Cell-Substrate Protrusions (CSPs). Similar to TMTs, CSPs generally originate from the lateral cell surface, contacting the substratum at their distal tips (Figure 1I, Supplementary Movie 2). While morphologically similar to TMTs, CSPs are slightly longer (Figure 1C) and wider (Figure 1D), although there is considerable heterogeneity in both parameters. The distal tips of CSPs tend to display an expanded width compared to the CSP shaft (Figure 1H). CSPs are 7.7-fold less abundant than TMTs (Figure 1B, mean 0.041±0.01 CSP/cell).

These experiments were conducted on cells plated onto fibronectin. We asked whether the nature of the substratum influenced the size or abundance of TMTs or CSPs. Plating onto the following coated surfaces results in minor variation of these parameters from fibronectin: uncoated glass, laminin, type-1 collagen, concanavalin A (Supplementary Figure 1).

### Cytoskeletal organization in DHPC-018 TMTs

We next examined the cytoskeletal components present in TMTs, initially focusing on actin filaments and microtubules. Using low-magnification fixed-cell imaging of actin filaments (TRITC-phalloidin), microtubules (anti-alpha-tubulin) and cell membranes (wheat germ agglutinin, WGA), we find that all TMTs contain actin filaments, and ∼93% contain microtubules (Figure 2A, B). We used higher resolution Airyscan microscopy to examine cytoskeletal organization in more detail. For TMTs of median thickness (∼2 μm), actin filament staining exists mainly along the cortex of the TMT, whereas microtubules enrich in the central region (Figure 2C, Supplementary Figure 2A). This organization of actin at the cortex is also observed upon 3D reconstruction of Z-stacks (Figure 2D). For thinner TMTs (<1 μm), the spatial organization is more difficult to determine at this resolution (Figure 2E, Supplementary Figure 2B).

**Fig. 2.**
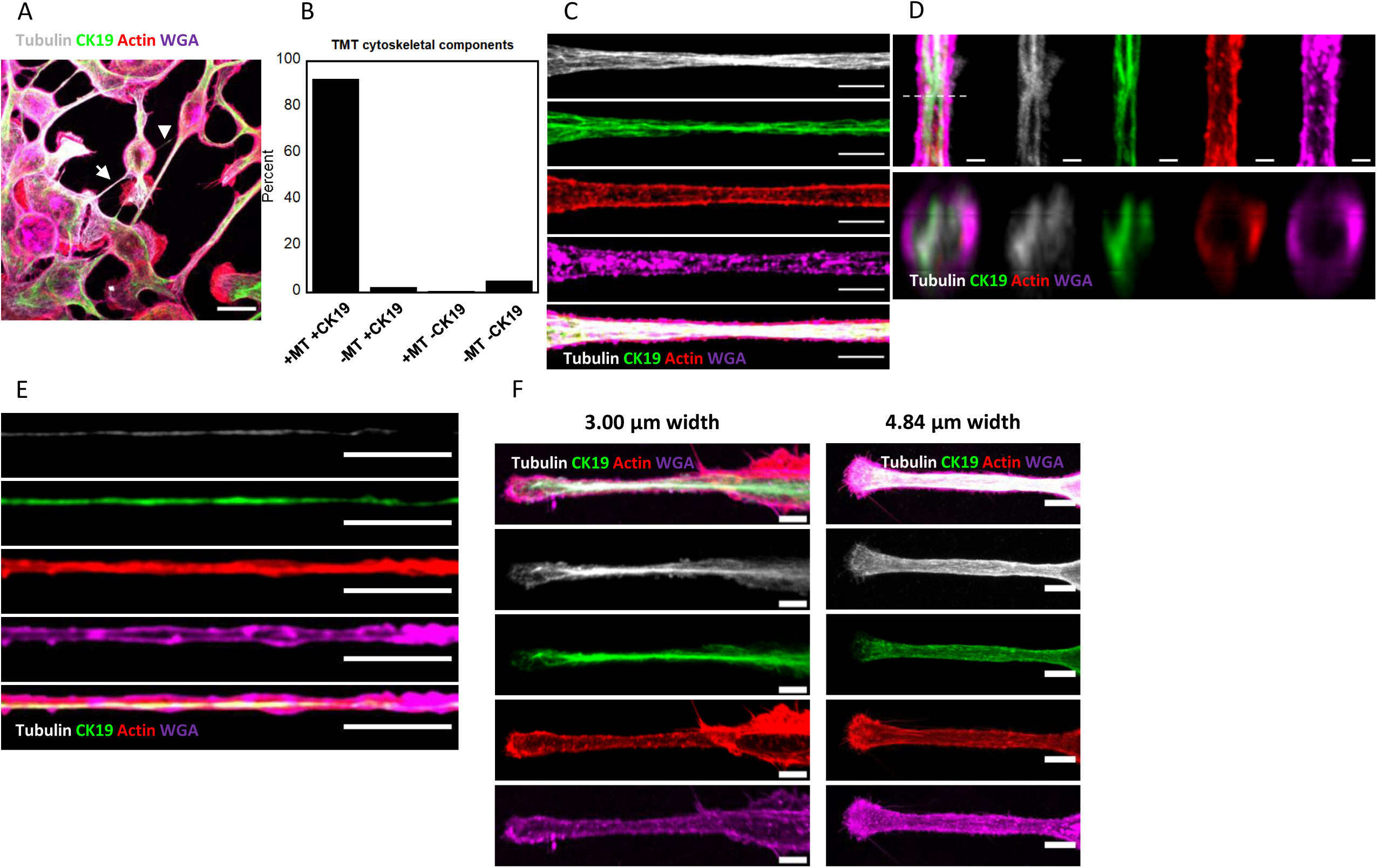
Cytoskeletal organization in DHPC-018 TMTs. Airyscan confocal images of cells plated on fibronectin, fixed and stained with anti-CK19 (green), anti-tubulin (white), TRITC-phalloidin (red), and WGA (magenta). (A) Max projection of a field of cells. Arrow indicates a TMT with actin, tubulin, and CK19; arrowhead indicates a TMT with only actin and CK19. Scale bar, 20 µm. (B) Percentage of TMTs containing CK19 and/or microtubules. All TMTs contain actin filaments. 611 TMTs. (C) High resolution image of a TMT of median thickness (∼2 μm). Scale bars, 5 µm. (D) 3D reconstruction of a TMT interior. Top: plan view. Scale bar, 1 µm. Bottom: XZ cross section of TMT, taken at the point indicated by the dashed line in the top panel. Actin filaments trace the TMT periphery, while CK19 and microtubules localize to the interior. (E) A TMT of thinner width (<1 μm). Scale bars, 5 µm. (F) Two examples of CSP cytoskeletal distribution. Left: CSP of intermediate width (average 3.00 μm) and length (44.0 μm). Right: CSP of larger width (average 4.84) length (70.5 μm). Scale bars, 5 µm.

We also examined DHPC-018 cells for the presence of cytokeratin 19 (CK19), since this intermediate filament protein has been used as a marker for PDAC (Jain *et al.*, 2010; Cen *et al.*, 2017). CK19 staining is enriched in ∼94% of TMTs (Figure 2A, B). Similar to microtubules, CK19-containing intermediate filaments (IFs) run through the interior of the TMT (Figure 2C, Supplementary Figure 2A). However, while IFs and microtubules are often found in close proximity in the TMT, they are not completely overlapping in localization, suggesting that they are not obligatorily associated.

Finally, we examined the cytoskeletal organization in CSPs. CSPs of two different widths are shown in Figure 2F, with additional examples in Supplementary Figure 2C. In the CSP shaft, organization is largely similar to TMTs, with actin enriched along the cortex while microtubules and CK19 are largely in the central region. For wider CSPs, there is some evidence for long filaments/bundles in the central region (Figure 2F, right). The CSP tip is more variable. In some cases, the tip is considerably wider than the shaft, with leading edge actin staining reminiscent of a lamellipodium or growth cone (Figure 2F). In other cases, the CSP tip is less broad, with greatly reduced leading edge actin staining (Supplementary Figure 2C).

### Assembly mechanisms and dynamics for DHPC-018 TMTs and CSPs

We used live-cell DIC microscopy to evaluate TMT assembly mechanisms and dynamics in DHPC-018 cells, allowing cells to adhere to fibronectin-coated coverslips for 4 hrs before acquiring images at multiple Z planes at 15- or 30-min intervals over the next 12-20 hrs. From these examinations, both TMTs and CSPs display a range of lifetimes, from one frame to over 12 hrs (Figure 3A).

**Fig. 3.**
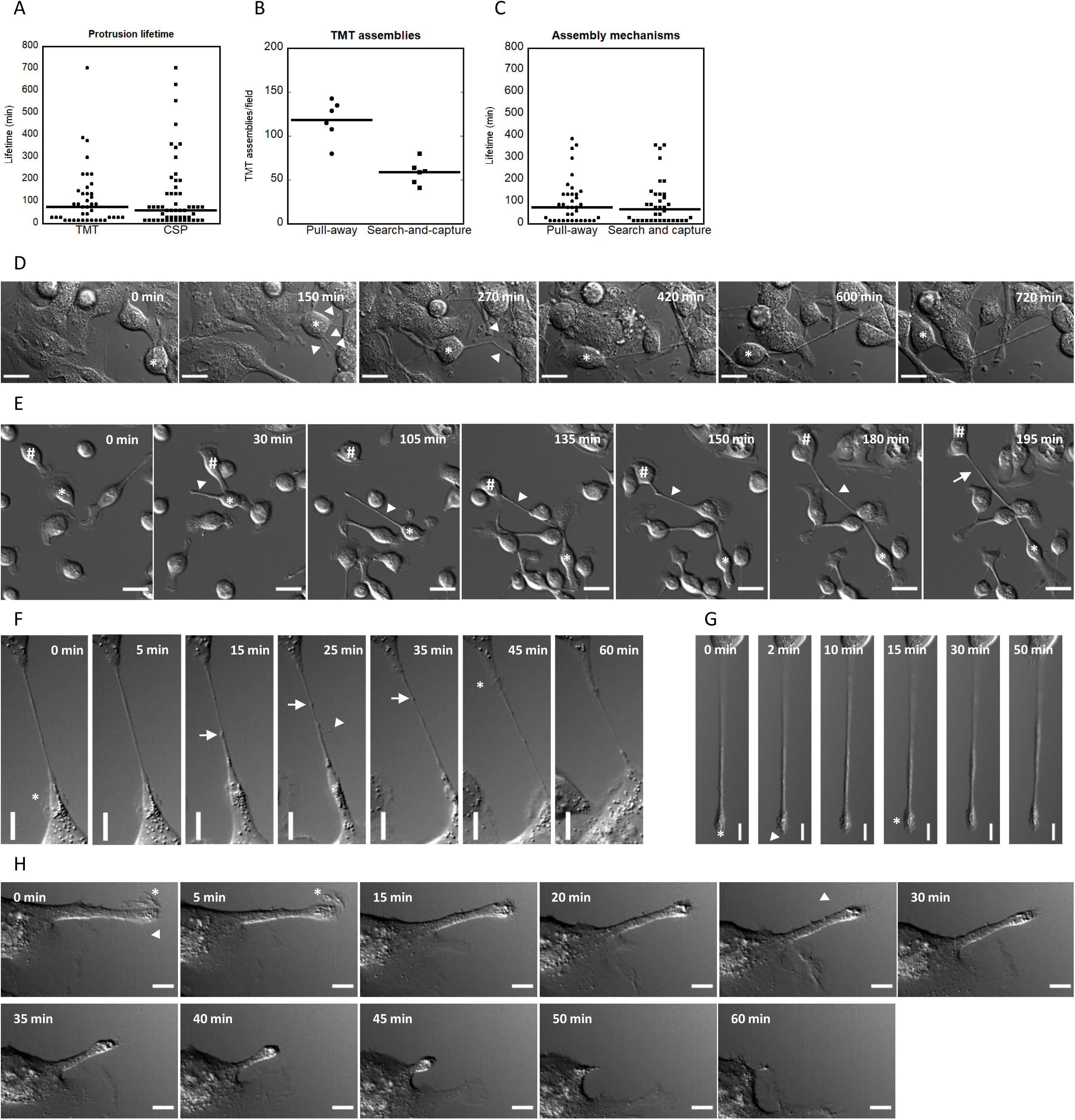
Assembly and dynamics of TMTs and CSPs. (A) Graph of protrusion lifetimes. Bars are medians, 75 ± 21.2 min for TMTs (40 events), 60 ± 22.4 min CSPs (52 events). (B) Graph of total TMT assemblies per field as a function of assembly mechanism. Bars are medians, 122 ± 9.29 for pull-away (710 events), 59.5 ± 5.51 for search-and-capture (352 events). (C) Graph of TMT lifetimes as a function of assembly mechanism. Bars are medians, 75 ± 17.5 min pull-away (35 events), 67.5 ± 17.2 min search- and-capture (35 events). (D) Time-lapse DIC montage (30 min frame interval) of pull-away assembly of a TMT. Asterisk depicts the cell that pulls away from the adjacent cells, creating three TMTs and a CSP (arrowheads) that appear to condense together. Z plane 3-4 µm above basal surface. Scale bar, 20 µm. From Supplementary Movie 3. (E) Time-lapse DIC montage (15 min frame interval) of search-and-capture TMT assembly. The cell marked with the asterisk extends a CSP (arrowhead) by moving away from the CSP tip. Another cell (marked by hashtag) makes contact with the CSP, then pulls away. The resulting TMT is denoted by an arrow. Scale bar, 20 µm. From Supplementary Movie 5. (F) Time-lapse DIC montage (30 sec frame interval) of an individual TMT, which undergoes periodic ruffling (asterisks), filopodial assembly (arrowhead), and translocation of a bulged region (arrows). Scale bar, 10 µm. Z plane 3-4 μm above basal surface. From Supplementary Movie 6. (G) Time-lapse DIC montage (30 sec frame interval) of an individual CSP, whose dynamic activity includes a filopodial assembly (arrowhead), ruffling (asterisks) and blebbing. Scale bar, 10 µm. Z plane on basal surface. From Supplementary Movie 7. (H) Time-lapse DIC montage (30 sec frame interval) of an individual CSP that undergoes retraction, preceded by termination of ruffling at the tip. Scale bar, 10 µm. Z plane on basal surface. From Supplementary Movie 8. Cells plated on glass coverslips in panels C and D, and on fibronectin-coated coverslips in panels E-G. All error calculations are SEM.

Similar to past studies (Sowinski *et al.*, 2008; Wang *et al.*, 2010; Gerdes *et al.*, 2013; Kumar *et al.*, 2017), we observe two main mechanisms of TMT assembly, which we term “pull-away” and “search-and-capture.” Pull-away represents ∼2/3 of the events under the plating conditions used here (Figure 3B). The lifetimes of TMTs created by either mechanism are similar (Figure 3C). For both mechanisms, a number of features suggest considerable flexibility in the process.

In pull-away assembly, one cell migrates away from another, leaving a TMT tether that can persist for multiple hours. Intriguingly, in some cases several TMTs can assemble between two cells in the early stages of pull-away, and subsequently condense into an apparent single TMT (Figure 3D, Supplementary Movie 3). In other cases, a single TMT is pulled between cells (Supplementary Figure 3A, Supplementary Movie 4).

In search-and-capture assembly, one cell extends a CSP and eventually makes contact with another cell (the ‘receiving’ cell). An interesting feature of search-and-capture in DHPC-018 cells is that it is not the CSP-containing cell that is doing the searching. CSPs are generally not elongating from their tips, but by the cell body moving away from the stationary CSP tip. In the context of search-and-capture, the receiving cell is the one that contacts the stationary CSP (Figure 3E, Supplementary Movie 5). In the example shown, the receiving cell then pulls away from the CSP, creating its own CSP in the process and resulting in a TMT containing segments from both cells.

After assembly, the TMT can undergo extensive changes in length, as well as being able to withstand considerable deformation by intervening cells (Figure 3D, E, Supplementary Movie 4, 5). We also examined TMT dynamics on a shorter timescale (60 min, 2 frames/min). In this context, additional dynamics are apparent including: protrusion of filopodia-like structures from the TMT; lamellipodia-like ruffling from the TMT/cell body interface; and directional movement of material along the TMT, suggestive of organellar contents (Figure 3F, Supplementary Movie 6). For CSPs, the tip does not generally translocate but displays considerable dynamics in the form of blebbing, filopodia or ruffling (Figure 3G, H, Supplementary Movie 7, 8). CSPs can undergo retraction, which is accompanied by a decrease in dynamics at the tip (Figure 3H, Supplementary Movie 8).

### TMT assembly is Arp2/3 complex-dependent

We next examined the roles of actin and microtubules in TMT assembly and maintenance, using latrunculin A (LatA, actin depolymerization) and nocodazole (Noc, microtubule depolymerization). Similar to past studies (Vallabhaneni *et al.*, 2012; Wittig *et al.*, 2012; Takahashi *et al.*, 2013; Han *et al.*, 2016), treatment with LatA for 24 hrs causes a significant decrease in TMTs, whereas TMT and CSP number increase with Noc treatment (Figure 4A-C). These differences are not due to effects on cell number, since neither LatA nor Noc affect this parameter significantly (Figure 4D). An additional effect occurring upon LatA treatment is the elaboration of numerous basal protrusions (Figure 4E). These protrusions are clearly distinct from TMTs, in that they are on the basal surface and are largely devoid of actin filaments, microtubules, and CK19. Similar basal protrusions occur upon use of another drug that affects actin depolymerization, cytochalasin D (Figure 4E). Since these protrusions occupy the basal plane, they preclude determination of CSP number with LatA treatment.

**Fig. 4.**
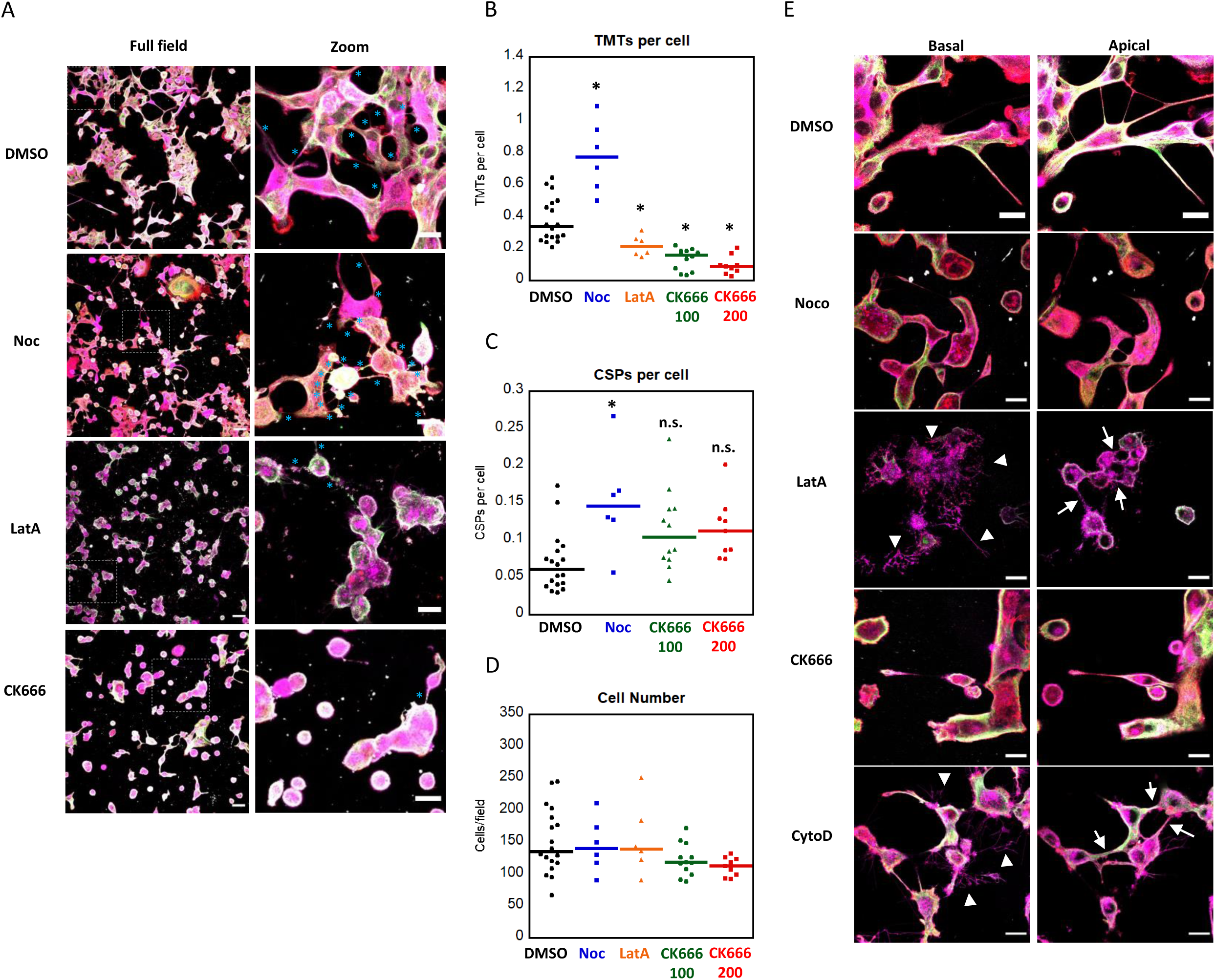
Actin depolymerization and Arp2/3 complex inhibition reduce TMT number. (A) Airyscan confocal max projections of fixed DHPC-018 cells following 24 hr treatment with DMSO, 50 µM Nocodazole, or 5 µM Latrunculin A. Boxed areas in the fields on the left correspond to the zooms on the right. Cells stained with anti-CK19 (green), anti-tubulin (white), TRITC-phalloidin (red), and WGA (magenta). Scale bars, 50 µm for full fields, 20 µm for zooms. (B) Number of TMTs per cell after 24 hr treatment. 5991 cells, 3606 TMTs total. 2725 cells, 1128 TMTs for DMSO; 872 cells, 720 TMTs for Noc; 927 cells, 210 TMTs for LatA; 1467 cells, 209 TMTs for 100 µM CK666; 1005 cells, 94 TMTs for 200 µM CK666. Bars are medians, 0.47 ± 0.06 DMSO, 0.77 ± 0.10 Noc, 0.21 ± 0.03 LatA, 0.16 ± 0.02 100 µM CK666, 0.09 ± 0.02 200 µM CK666. * indicates p ≤ 0.05 by ANOVA with Tukey’s Honest Significant Difference. (C) Number of CSPs per cell after 24 hr treatment with DMSO, 50 µM Nocodazole, or 200 µM CK666. 182 CSPs DMSO, 119 Noc, 156 100 µM CK666, 115 200 µM CK666. Bars are medians, 0.07 ± 0.01 DMSO, 0.15 ± 0.03 Noc, 0.10 ± 0.02 100 µM CK666, 0.11 ± 0.01 200 µM CK666. CSPs in LatA-treated cells were not quantified, due to the extensive basal protrusions formed. * indicates p ≤ 0.05 by ANOVA with Tukey’s Honest Significant Difference, n.s. indicates no statistical significance. (D) Number of cells per field after the indicated 24 hr treatment. Bars are medians, 135.5 ± 12.06 DMSO, 140 ± 17.46 Noc, 139.5 ± 22.96 LatA, 118.5 ± 7.31 100 µM CK666, 113 ± 4.92 200 µM CK666. (E) Airyscan confocal images of DHPC-018 cells fixed after DMSO, Latrunculin A, Cytochalasin D, Nocodazole, and CK666 treatments. Left: single 0.4 µm Z slice basal images, right: single Z slice apical images. Arrowheads indicate the basal retraction fibers following LatA and CytoD treatments, while arrows indicate apical TMTs. Staining as in panel A. Scale bars, 20 µm. All error calculations are SEM.

To probe the role of actin in more detail, we utilized an inhibitor of Arp2/3 complex, CK666. Arp2/3 complex is a major actin nucleation factor, and is required for a wide range of cellular actin-based structures (Campellone and Welch, 2010). Treatment with CK666 for 24 hrs causes a significant decrease in TMTs (Figure 4A,B) without causing a significant drop in cell number (Figure 4D). Unlike LatA, CK666 treatment does not result in basal surface protrusions (Figure 4E).

We also examined the effects of CK666 treatment on live cells, in order to determine the mechanism leading to TMT loss. Over a 3-hr treatment period, CK666-treated cells display a 66% decrease in TMT assembly events (Figure 5A). This decrease is consistent over the experiment time course (Figure 5B), suggesting that Arp2/3 complex is acutely required for TMT assembly. Both pull-away and search-and-capture events are reduced by this treatment (Figure 5C). CK666 treatment does not cause a dramatic change in cell morphology over this time course (Figure 5D), in contrast to the effects of LatA. Upon minutes after LatA treatment, cells retract to leave the basal protrusions, indicating that these apparent protrusions are actually retraction fibers (Figure 5D).

**Fig. 5.**
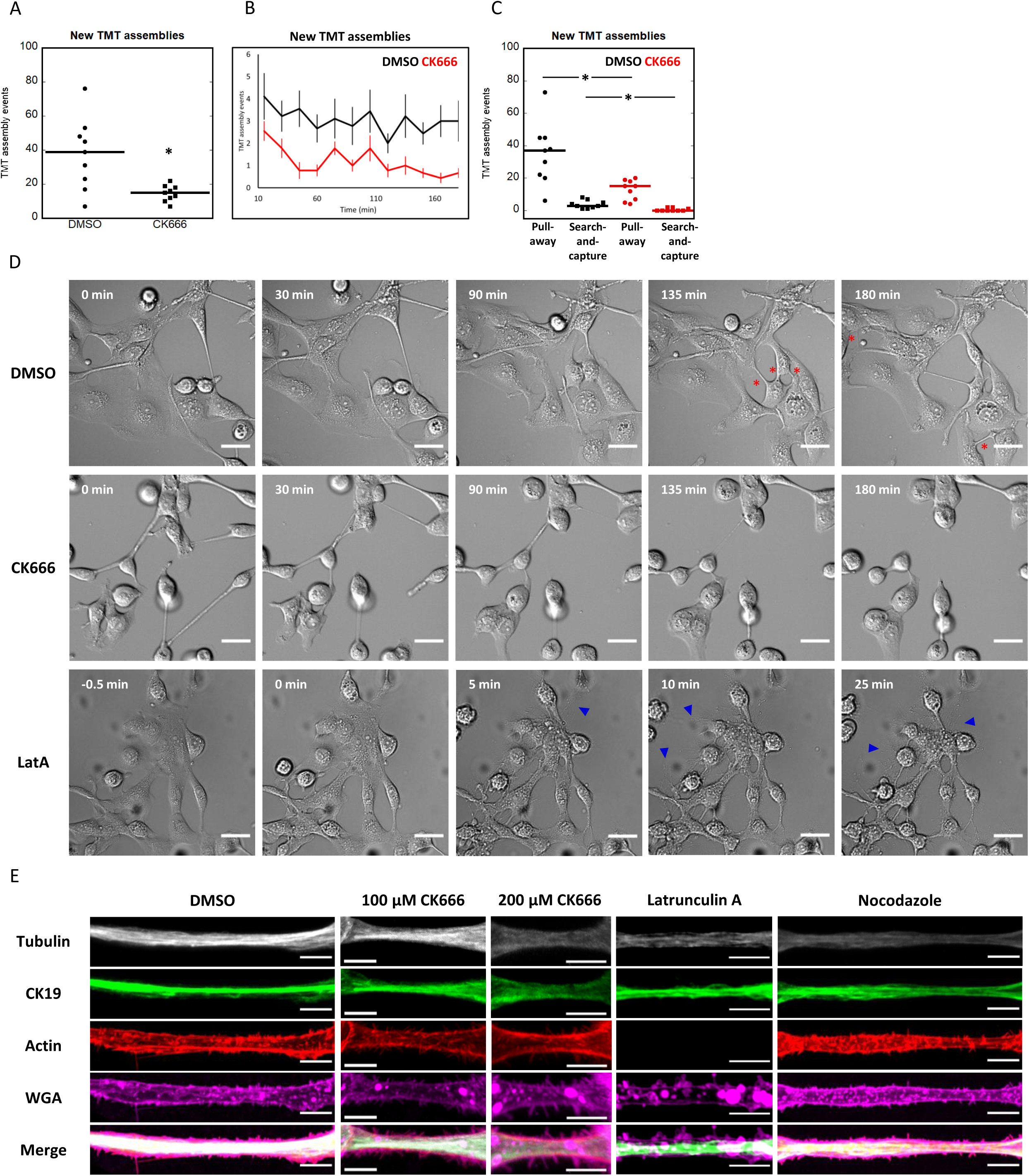
Arp2/3 acts in assembly of new TMTs. (A) Number of total TMT assembly events quantified from live DIC imaging over 3 hours of treatment with DMSO or 200 µM CK666. 350 DMSO TMT assemblies, 120 CK666 TMT assemblies. Bars are medians, 39 ± 6.97 DMSO, 15 ± 4.69 CK666. * indicates p < 0.005 by Wilcoxon Rank Sum test. (B) TMT assemblies as a function of time during DMSO or 200 µM CK666 treatments. (C) TMT assemblies by type following treatment with DMSO or 200 µM CK666. Bars are medians, 37 ± 6.36 pull-away and 3 ± 0.83 search-and-capture for DMSO, 15 ± 2.04 pull-away and 0 ± 0.30 search-and-capture for CK666. * indicates p < 0.005 by unpaired student T test. (D) Live DIC montages of DHPC-018 cells over 3 hr treatments with DMSO (top), 200 µM CK666 (middle) or 5 µM LatA (bottom). Red asterisks denote new TMT assembly. Blue arrowheads denote basal protrusions. Note that LatA montage is taken over a different time period. (E) Examples of TMTs that persist following 1 hr of treatment with DMSO, 5 µM LatA, 50 µM nocodazole, or 100 µM CK666. Cells stained with anti-CK19 (green), anti-tubulin (white), TRITC-phalloidin (red), and WGA (magenta). Scale bars, 5 µm. All error calculations are SEM.

Finally, we examined cytoskeletal distribution in TMTs after inhibitor treatments. Actin filament staining is largely eliminated from TMTs after a 1-hr treatment with LatA (Figure 5E). In contrast, actin filaments persist in CK666-treated cells over this time period (Figure 5E). Nocodazole treatment results in loss of microtubules, but actin filaments stay intact (Figure 5E). These results suggest that Arp2/3 complex is not required for continuous maintenance of actin filaments in existing TMTs.

### TMTs in PDAC tumors

To test whether TMTs are present in a tumor-like environment, we injected DHPC-018 cells into the flanks of immune-compromised mice, resulting in tumors of approximately one cm diameter in six weeks. Formalin-fixed, paraffin-embedded (FFPE) sections of the excised tumors were stained with anti-CK19 and DAPI. This approach resulted in a dense CK19 staining pattern, due to the fact that DHPC-018 cells represent close to 100% of the cells in the tumor (Supplementary Figure 4A). The dense cell packing precluded detection of TMTs connecting individual DHPC-018 cells.

As an alternative approach, we created a DHPC-018 cell line stably expressing GFP and raised flank tumors in which 10% of the injected cells were GFP-expressing. Low-resolution tile-scans reveal GFP throughout the tumor (Figure 6A). High-resolution imaging with 3D reconstruction reveals readily discernable TMTs, regardless of the fixation/preparation techniques used (Figure 6B, Supplementary movie 9). These results suggest that TMTs are maintained in DHPC-018 cells in a tumor-like environment.

**Fig. 6.**
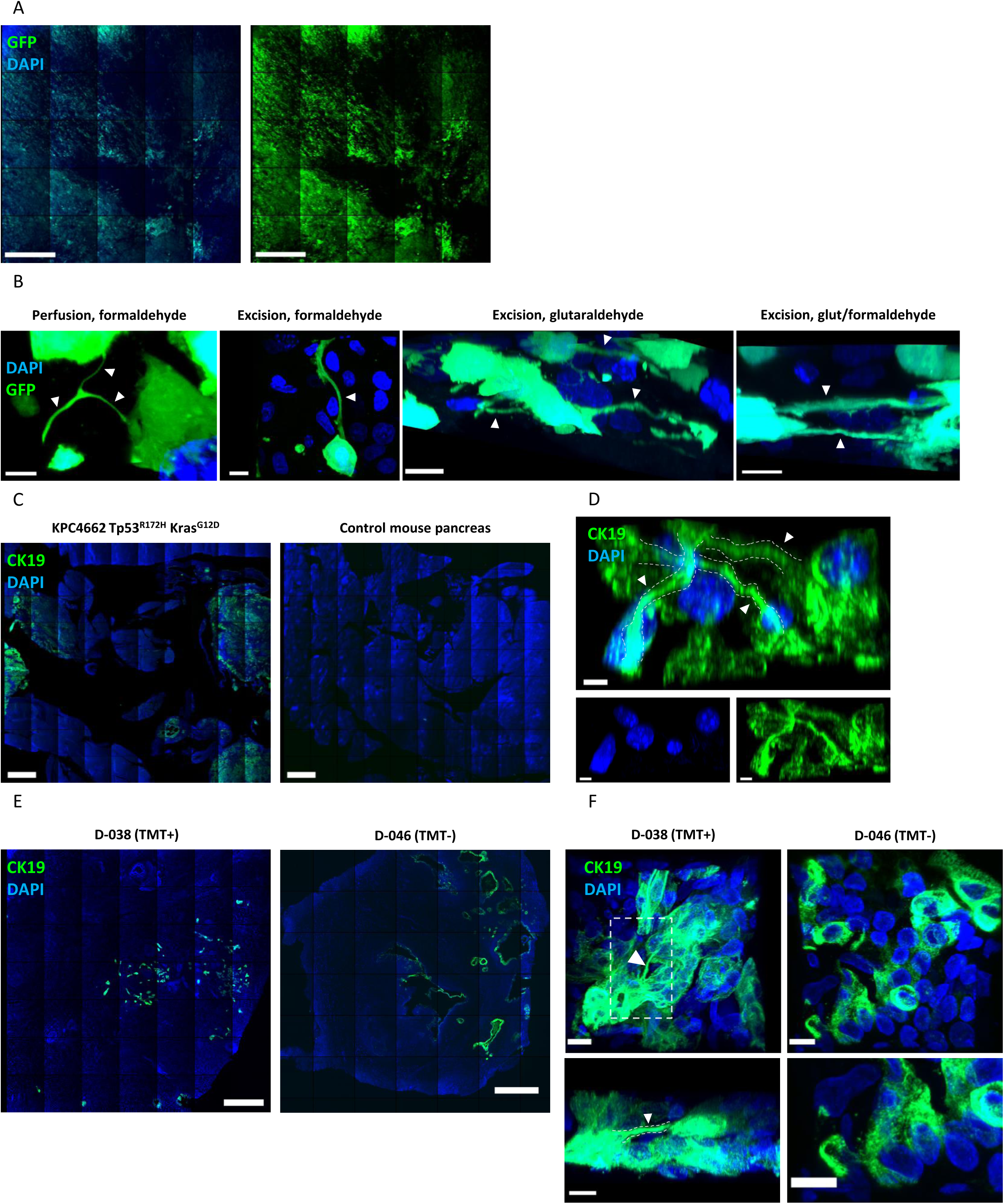
TMTs in the PDAC tumor environment. (A) Low magnification (10X objective) tile-scan image of a section from a DHPC-018 flank tumor, in which 10% of the cells express GFP. Section stained with DAPI (blue). Left: overlay, right: GFP alone, enhanced contrast. Scale bars, 500 µm. (B) Representative higher magnification (63x objective) 3D reconstruction images of four DHPC-018 flank tumors (10% GFP). Left to right: formaldehyde perfusion fixation by cardiac puncture, excision and fixation by formaldehyde, excision and fixation by glutaraldehyde, excision and fixation by combination of formaldehyde and glutaraldehyde (From Supplementary Movie 9). Arrowheads indicate GFP-positive TMTs. Scale bars, 10 μm for excision fixations, 5 μm for perfusion fixation. (C) Low magnification (10X objective) tile-scan images of FFPE pancreas sections from C57BL/6 mice. Left: pancreas after tumor formation by orthotopically-injected KPC4662 acinar cells. Right: control pancreas. Sections stained with DAPI (blue) and anti-CK19 (green). Scale bars, 500 µm. (D) Higher magnification (63x objective) 3D reconstruction image of the KPC4662 tumor from C. Arrowheads/outlines indicate CK19^+^ TMTs. From Supplementary Movie 10. Scale bars, 5 µm. (E) Representative low magnification tile-scans of sections from FFPE samples of primary tumors from two patients. Sections stained with DAPI (blue), and anti-CK19 (green). Scale bars, 500 µm. (F) Higher magnification (63x objective) 3D reconstruction images of the tumors from E, focused on CK19^+^ regions. Arrowheads/outlines indicate CK19^+^ TMTs in tumor D-038, while tumor D-046 appears negative for CK19^+^ TMTs. Bottom left: XZ view of TMT boxed in D-038, arrowhead indicates TMT. Bottom right: closer view of CK19+ region of D-046, devoid of TMTs. Sections stained with DAPI (blue), and anti-CK19 (green). From Supplementary Movie 11. Scale bars, 10 µm.

We next extended our study to a mouse model system in which PDAC is established in an immune-competent environment by injection of cultured KPC 4662 (Kras^G12D/+^, Trp53^R172H/+^, Pdx1-Cre) pancreatic acinar cells into the pancreas of C57BL/6 mice (Winograd *et al.*, 2015). The low cellularity of these tumors allows evaluation of TMTs by CK19 staining. Tile-scans reveal elevated and localized CK19 staining compared to control mouse pancreas (Figure 6C). High resolution imaging with 3D reconstruction reveals CK19-positive TMT connections between cells (Figure 6D, Supplementary Movie 10).

To assess the presence of TMTs in a clinically-relevant situation, we imaged CK19-stained FFPE sections of six PDAC tumors from Dartmouth Hitchcock Medical Center (Table 1). Five samples were resected tumors (four from pancreas, one from a liver metastasis), and one was a needle biopsy core. All patients had undergone neoadjuvant treatment prior to resection/biopsy. Low-resolution tile-scans revealed sporadic pockets of CK19-positive cells (Figure 6E), which were more closely examined at higher resolution for TMTs. We identified TMTs in three tumors, while two tumors displayed no detectable TMTs by our analysis (Figure 6F, Supplementary Movie 11).

## Discussion

In this study, we identify a population of inter-cellular linear protrusions in a pancreatic cancer (PDAC) cell line, which we define as tumor microtubes (TMTs). In addition to these structures, a second type of protrusion of similar morphology makes contact with the substratum rather than with another cell, and we define these as cell-substrate protrusions (CSPs). Both TMTs and CSPs emanate from the lateral plasma membrane as opposed to the basal surface, a property similar to TNTs (Rustom *et al.*, 2004; Gurke *et al.*, 2008; Abounit and Zurzolo, 2012). The majority of TMTs and CSPs contain three cytoskeletal elements: actin filaments, microtubules, and intermediate filaments (IFs), with actin filaments being enriched along the membrane while microtubules and IFs run along the center. Live-cell microscopy shows that two assembly mechanisms are employed for TMTs, which we term “pull-away” and “search-and-capture” and are similar to previously elucidated TNT assembly mechanisms (Sowinski *et al.*, 2008; Veranič *et al.*, 2008; Gerdes *et al.*, 2013; Kumar *et al.*, 2017). Actin polymerization through Arp2/3 complex is necessary for TMT assembly. We observe TMTs in PDAC tumors in a number of contexts, including a sub-set of primary human tumors.

An important finding of our work is that the structures we identified are quite heterogeneous in a number of characteristics, including width and assembly mechanism. For this reason, we struggled with which name to assign to the structures. We ultimately chose TMTs for the following reasons: the majority are in the larger width range (>1 μm), they are present in a tumor-derived cell line, and they contain microtubules. However, 21% of the structures are <1 μm and 9% are < 0.5 μm in width, which is in the range broadly defined for TNTs. The heterogeneity in this population suggests the possibility of fundamentally distinct functions for these structures. A second point to consider is that the width of an individual TMT can vary over time, which is apparent from our live-cell DIC imaging. A previous study of TNTs from neuronal cell lines showed that multiple TNTs could bundle together into a larger structure (Sartori-Rupp *et al.*, 2019). This scenario is unlikely here, due to the absence of plasma membrane staining within the thicker TMTs we observe. In addition, the structures observed in neuronal cell lines were devoid of microtubules, which is not the case here.

Another heterogeneous aspect of this TMT population is in assembly mechanism. While we broadly use two assembly categories that have been previously defined for TNTs (Sowinski *et al.*, 2008; Veranič *et al.*, 2008; Gerdes *et al.*, 2013; Kumar *et al.*, 2017), our live-cell imaging results suggest heterogeneity within both mechanisms. For the pull-away mechanism, there is a sub-set of cases in which multiple TMTs assemble initially and then zipper into one. In the search-and-capture mechanism, the ‘receiving’ cell can pull away from the ‘donor’ cell (the cell extending the CSP), creating a hybrid TMT containing material from both cells. Such hybrid structures have been observed previously for TNTs (Sowinski *et al.*, 2008). Another interesting feature of search-and-capture is that the receiving cell appears to be the one to initiate contact, in that it migrates to the comparatively immobile CSP. This process is fundamentally different from TNT search-and-capture observed in PC12 (Rustom *et al.*, 2004), immune cells (Önfelt and Davis, 2004), or cytonemes (Ramírez-Weber and Kornberg, 1999), in which filopodia protrude to make contact with the receiving cell.

The organization of cytoskeletal elements within TMTs in our system was surprising to us. The most detailed examination of actin organization in TNTs shows long, parallel filaments akin to those in filopodia and microvilli (Sartori-Rupp *et al.*, 2019), although the actin-binding proteins involved appear to be different for filopodia and TNTs (Delage *et al.*, 2016). In the case of PDAC TMTs, actin clearly enriches along the plasma membrane, with little evidence of filaments running down the center of the structure. Instead, both microtubules and intermediate filaments occupy the central region. These findings suggest that neither TMTs nor CSPs are an elaboration of filopodia, but rather are more similar to neuronal-like processes, as suggested for cytonemes (Roy *et al.*, 2014; Huang *et al.*, 2019).

Actin filament localization near the plasma membrane is reminiscent of the cortical actin found in the cell body of many cell types (Chugh *et al.*, 2017). Cortical actin consists of a meshwork of actin filaments and myosin II, and the contractile force exerted by the myosin resists cellular turgor pressure. Arp2/3 complex has been shown to be an important assembly factor for cortical actin (Bovellan *et al.*, 2014). It is, therefore, surprising that the actin filament staining in TMTs is maintained after a 1 hr treatment with the Arp2/3 complex inhibitor CK666, whereas a 1 hr treatment with the actin sequestering molecule LatA results in loss of actin staining.

Our results show, however, that Arp2/3 complex is important in TMT assembly. The exact mechanism by which Arp2/3 complex participates in TMT assembly remains to be defined. Both pull-away and search-and-capture involve translocation of the entire cell, generally considered to involve Arp2/3 complex (Blanchoin *et al.*, 2014). However, cell-cell adhesion also involves Arp2/3 complex (Collins *et al.*, 2017), so TMT assembly could also be affected at this level.

TMTs in our DHPC-018 cells also contain microtubules and intermediate filaments. While microtubules are absent from some TNTs (Rustom *et al.*, 2004; Sowinski *et al.*, 2008; Wang *et al.*, 2010; Desir *et al.*, 2018; Kretschmer *et al.*, 2019), they are present in others (Önfelt *et al.*, 2006; Osswald *et al.*, 2015; Jansens *et al.*, 2017; Sáenz-de-Santa-María *et al.*, 2017; Resnik *et al.*, 2018; Zhang *et al.*, 2019). Where examined, the microtubule-containing structures are wider than those lacking microtubules (Önfelt *et al.*, 2006). Microtubules could play clear roles in organelle trafficking within TMTs, but also may be important in maintaining the wider diameter of TMTs (Zhang *et al.*, 2019). A number of intermediate filament proteins have been identified in TNTs, including cytokeratins and vimentin (Veranič *et al.*, 2008; Ady *et al.*, 2014; Sáenz-de-Santa-María *et al.*, 2017; Resnik *et al.*, 2018). We find abundant CK19 in TMTs from pancreatic cancer cells both in culture and in a tumor environment. Given the role of keratin-containing intermediate filaments in resisting mechanical tension in epithelial cells (Coulombe and Omary, 2002), CK19 may serve to resist the significant tension exerted on TMTs during their lifetime (see for example Supplementary Movie 3). It is possible that the presence of CK19 and other intermediate filament proteins allow TMTs to resist fixation by formaldehyde, (this paper and unpublished observations), while thinner TNTs are much more labile to fixation (Rustom *et al.*, 2004; Koyanagi *et al.*, 2005; Watkins and Salter, 2005).

An interesting question relates to the specific composition of the intermediate filaments in TMTs, given the ∼70 filament forming intermediate filament proteins found in mammals (Chung *et al.*, 2013). Thus far, CK7 (Veranič *et al.*, 2008; Resnik *et al.*, 2018), CK19 (this study), and vimentin (Ady *et al.*, 2014; Sáenz-de-Santa-María *et al.*, 2017) have been identified in this context. In view of the growing appreciation for the pleiotropic roles of intermediate filament in cancer (Karantza, 2011), their function in TNTs and TMTs assumes greater importance.

Our finding that TMTs are present in the pancreatic tumor environment agrees with previous observations in several tumor types (Lou *et al.*, 2012; Osswald *et al.*, 2015; Griessinger *et al.*, 2017; Desir *et al.*, 2018). One function of TMTs is to provide resistance to radiation, chemotherapy, or other assaults (Osswald *et al.*, 2015; Weil *et al.*, 2017; Desir *et al.*, 2018). Interestingly, we can only identify clear TMTs in a sub-set of primary tumor samples. Expansion of this study to a larger number of tumors, and development of additional TMT markers, is needed to determine the features of TMT-containing tumors in terms of TMT initiation and role in tumor progression. The differential presence of TMTs could provide an additional means of stratification of pancreatic cancer, which would inform treatment and eventual use of TMT-disrupting therapies.

---

**Supplementary Table 1.**
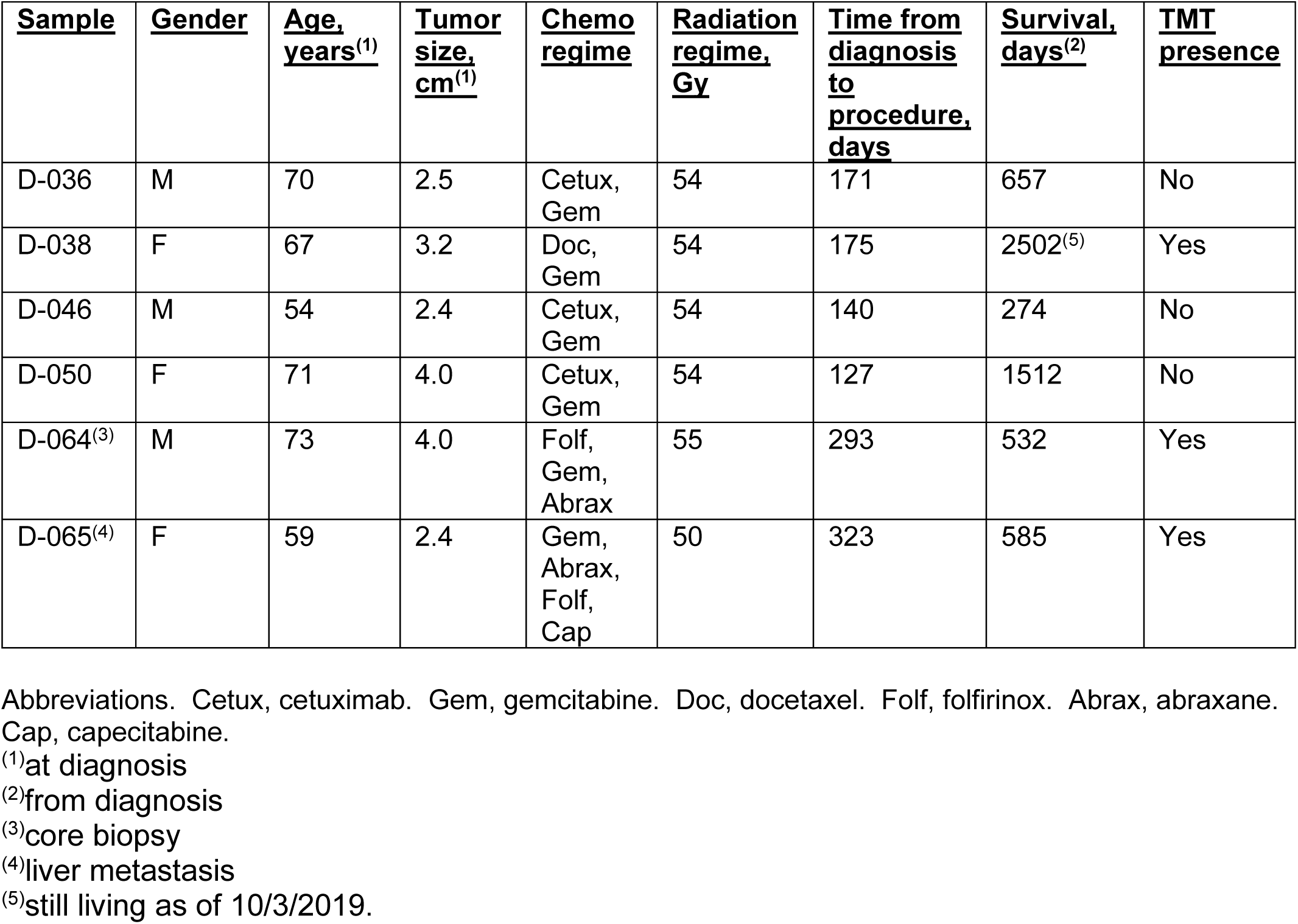
Human tumor samples.

| <u>Sample</u> | <u>Gender</u> | <u>Age, years<sup>(1)</sup></u> | <u>Tumor size, cm<sup>(1)</sup></u> | <u>Chemo regime</u> | <u>Radiation regime, Gy</u> | <u>Time from diagnosis to procedure, days</u> | <u>Survival, days<sup>(2)</sup></u> | <u>TMT presence</u> |
| --- | --- | --- | --- | --- | --- | --- | --- | --- |
| D-036 | M | 70 | 2.5 | Cetux, Gem | 54 | 171 | 657 | No |
| D-038 | F | 67 | 3.2 | Doc, Gem | 54 | 175 | 2502<sup>(5)</sup> | Yes |
| D-046 | M | 54 | 2.4 | Cetux, Gem | 54 | 140 | 274 | No |
| D-050 | F | 71 | 4.0 | Cetux, Gem | 54 | 127 | 1512 | No |
| D-064<sup>(3)</sup> | M | 73 | 4.0 | Folf, Gem, Abrax | 55 | 293 | 532 | Yes |
| D-065<sup>(4)</sup> | F | 59 | 2.4 | Gem, Abrax, Folf, Cap | 50 | 323 | 585 | Yes |

Abbreviations. Cetux, cetuximab. Gem, gemcitabine. Doc, docetaxel. Folf, folfirinox. Abrax, abraxane. Cap, capecitabine.
<sup>(1)</sup>at diagnosis
<sup>(2)</sup>from diagnosis
<sup>(3)</sup>core biopsy
<sup>(4)</sup>liver metastasis
<sup>(5)</sup>still living as of 10/3/2019.

**Supplementary Figure 1.**
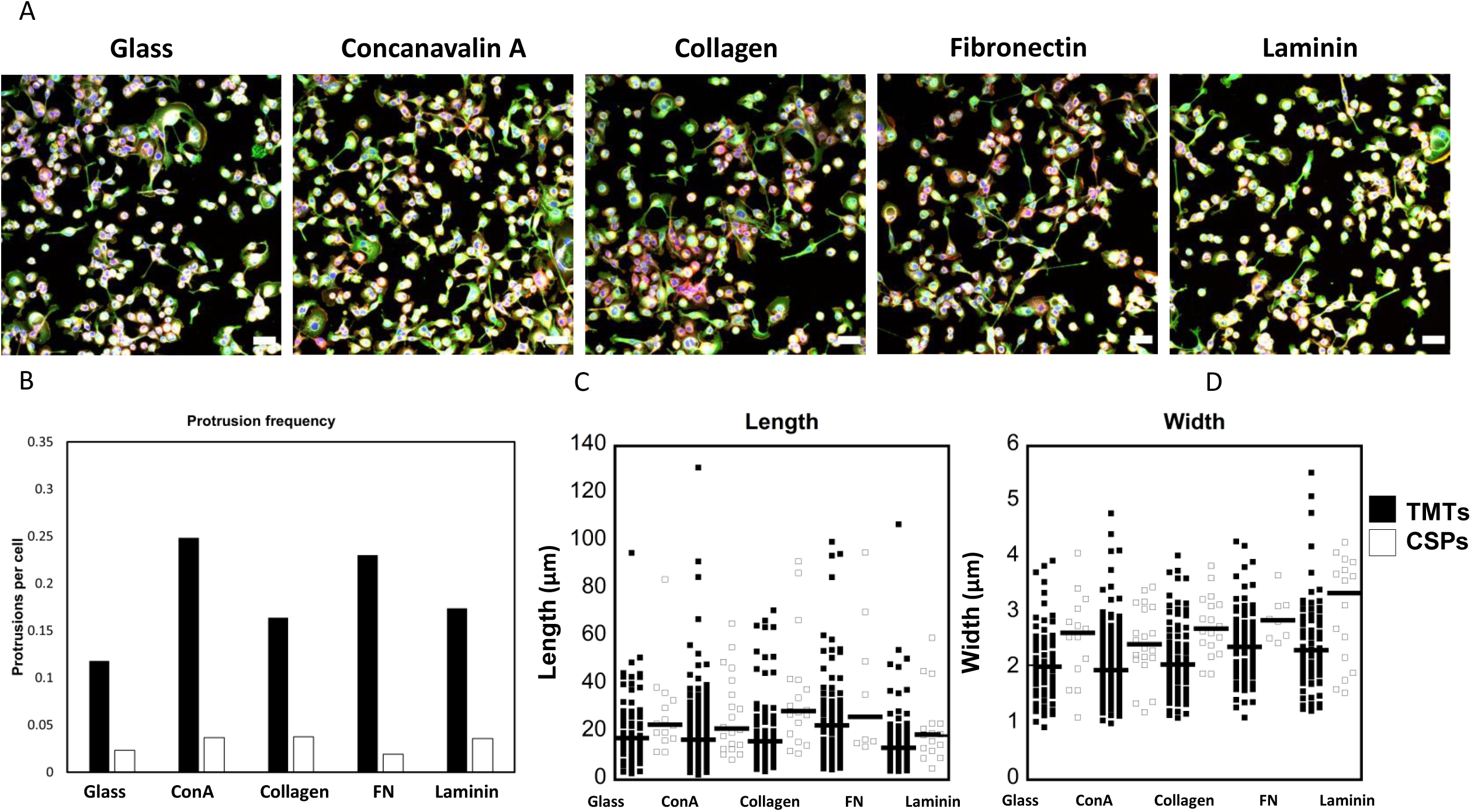
DHPC-018 TMT and CSP characteristics on different substrata. Cells were plated onto the indicated substrata for 16 hrs, then fixed and stained with DAPI (blue), mitochondria (gray), CK19 (green), and actin filaments (red). (A) Representative cell fields taken with the 20x 0.75 NA objective. Maximum intensity projections of 0.5 μm slices (24-33 slices). Scale bar, 50 μm. (B) TMT and CSP number per cell. 1902 cells, 451 TMTs, 76 CSPs total. 603 cells, 71 TMTs, 14 CSPs for glass; 548 cells, 136 TMTs, 20 CSPs for ConA; 483 cells, 79 TMTs, 18 CSPs for Collagen; 422 cells, 97 TMTs, 8 CSPs for FN; 449 cells, 78 TMTs, 16 CSPs for Laminin. (C) TMT and CSP length. Bars are medians ± standard error of the mean for TMTs and CSPs, respectively: 17.46 ± 1.75 and 23.03 ± 4.88 μm (glass), 16.77 ± 1.47 and 21.45 ± 3.72 (ConA), 16.10 ± 1.81 and 28.76 ± 5.56 (collagen), 22.75 ± 1.98 and 26.36 ± 10.76 (fibronectin), 13.43 ± 1.73 and 18.94 ± 3.76 (laminin). (D) TMT and CSP widths. Medians ± standard error of the mean for TMTs and CSPs, respectively: 2.03 ± 0.08 and 2.64 ± 0.21 μm (glass), 1.97 ± 0.06 and 2.43 ± 0.14 (ConA), 2.07 ± 0.08 and 2.71 ± 0.13 (collagen), 2.39 ± 0.06 and 2.87 ± 0.14 (fibronectin), 2.33 ± 0.09 and 3.36 ± 0.24 (laminin).

**Supplementary fig. 2.**
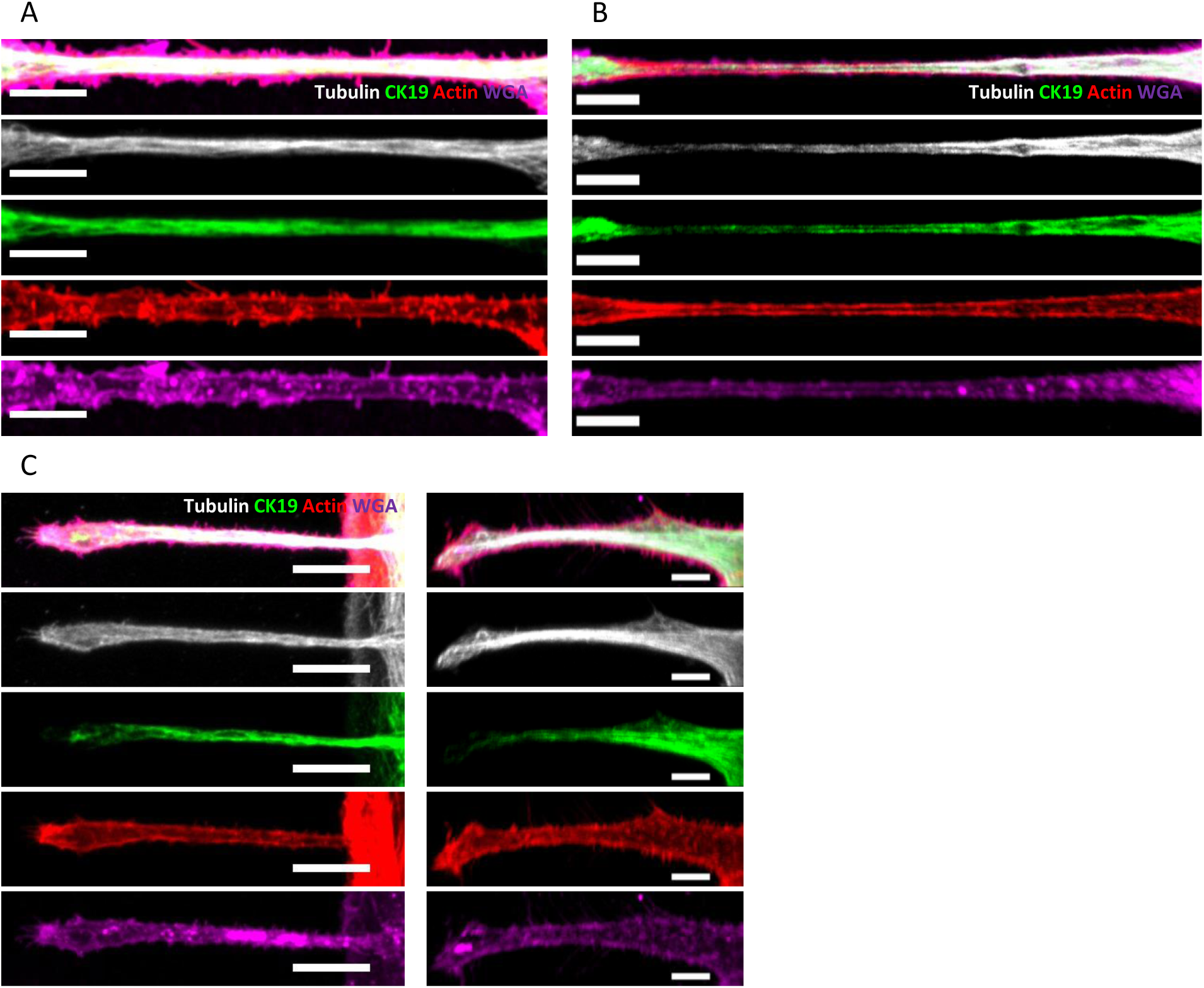
Further examples of TMT and CSP cytoskeletal distribution. Airyscan confocal images of DHPC-018 cells plated on fibronectin, fixed and stained with anti-CK19 (green), anti-tubulin (white), TRITC-phalloidin (red), and WGA (magenta). (A, B) TMTs. (A) shows a TMT of average length (35.5 μm) and width (1.80 μm average), while (B) is longer (43.1 μm) and thinner (0.65 μm average). Scale bars, 5 µm. (C) CSPs with average widths of 2.01 (left) and 3.41 μm (right); and lengths of 44.9 (left) and 42.3 μm (right). Scale bars, 10 µm (left) and 5 µm (right).

**Supplementary fig. 3.**
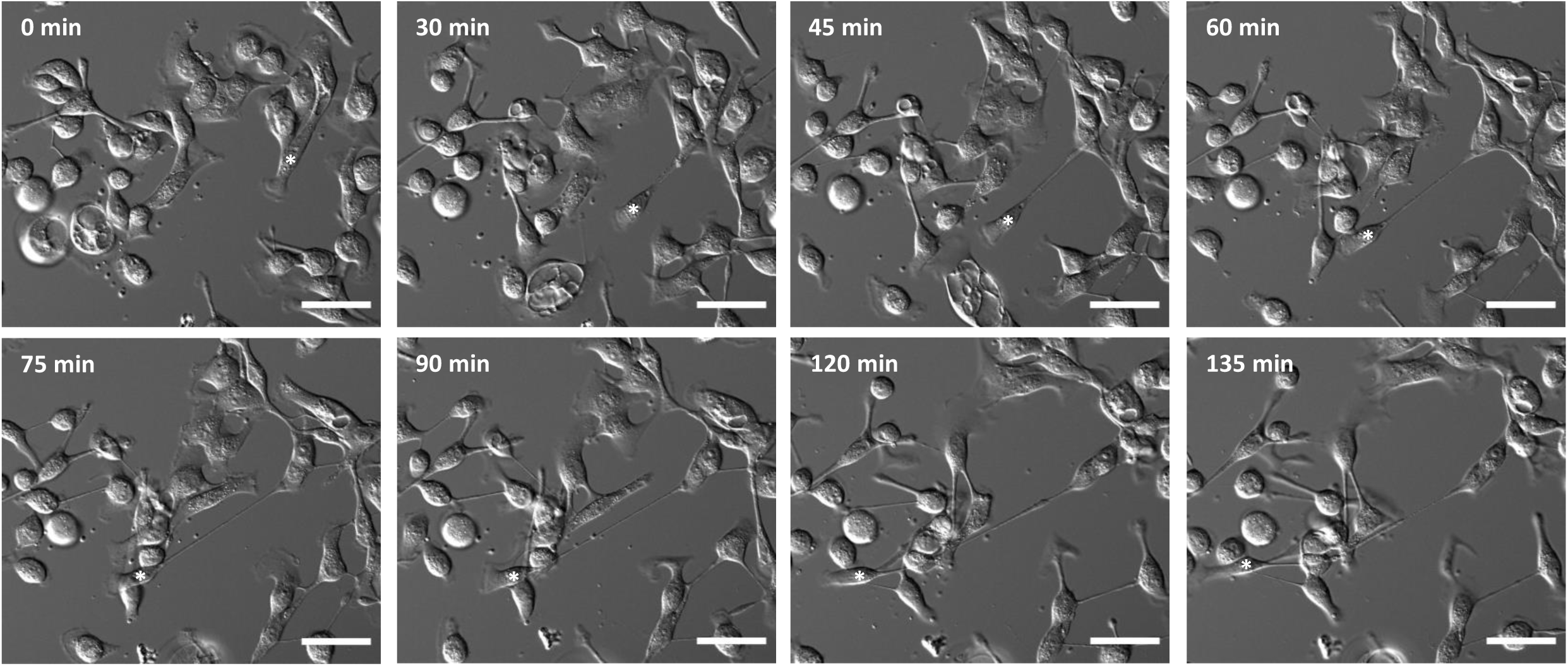
Pull-away TMT assembly. Time-lapse montage of pull-away assembly of a single TMT. DIC imaging of DHPC-018 cells on a glass coverslip. Asterisk depicts the cell that pulls away from the adjacent cells, creating one long TMT. Z plane 2-3 µm above basal surface. Scale bar, 50 µm. From Supplementary Movie 4.

**Supplementary fig. 4.**
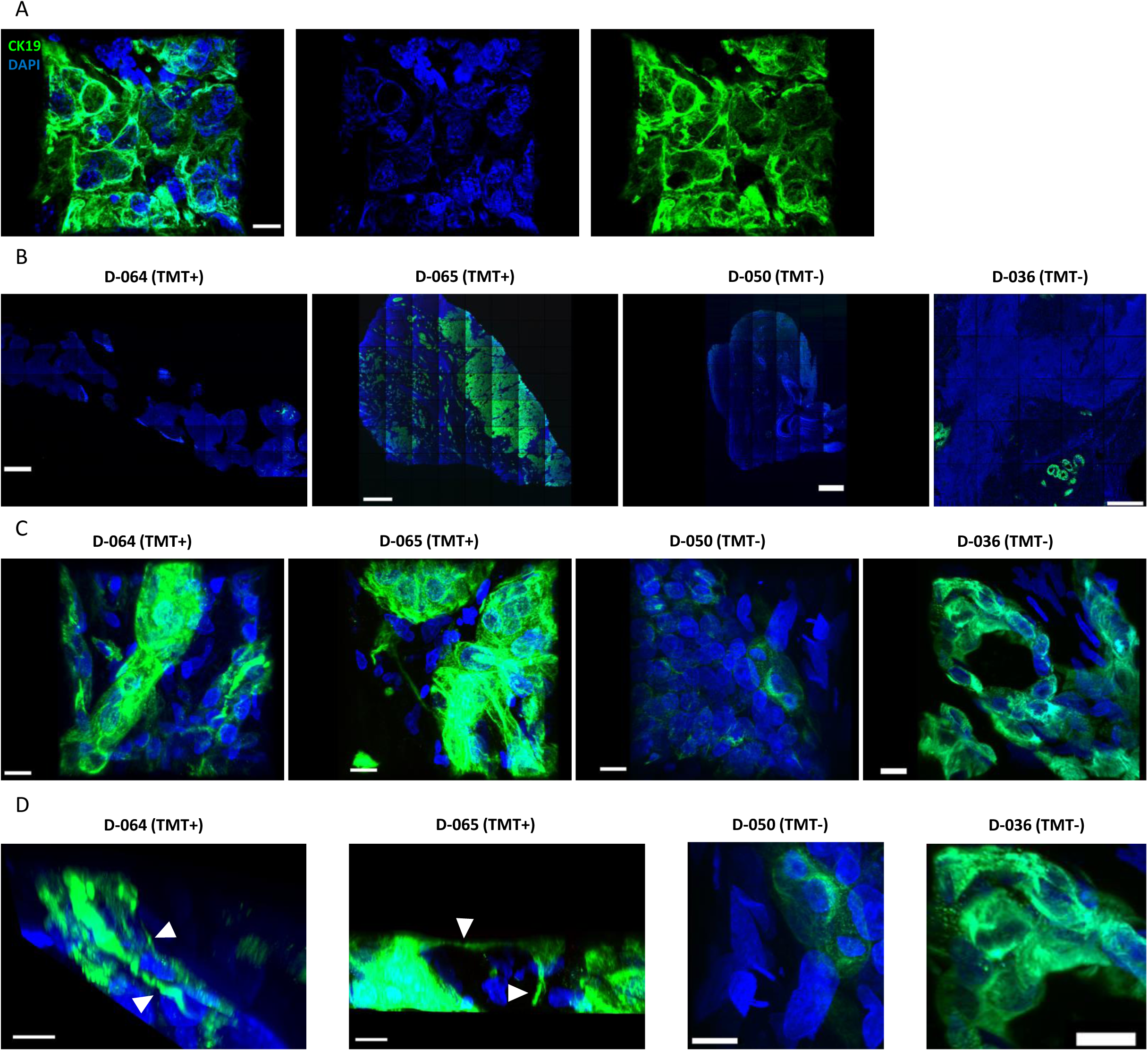
TMTs in the tumor environment. (A) DHPC-018 flank tumor grown in an immune-compromised NSG mouse. FFPE sample sectioned and stained with DAPI (blue) and anti-CK19 (green). Imaged on Zeiss LSM 880 Airyscan (63x). Scale bars, 10 µm. (B-D) FFPE sections of primary tumor samples, stained with DAPI (blue) and anti-CK19 (green). Imaged on Zeiss LSM 880 Airyscan. (B) 10x or 20x tilescan images. Scale bars, 500 µm D-064 (10x), 500 µm D-065 (10x), 500 µm D-050 (10x), 200 µm D-036 (20x). (C) 63x images of 3D reconstructions. Scale bars, 10 µm. (D) Closer views of images from C, re-oriented in 3D space. Arrowheads indicate TMTs in D-064 and D-065. Scale bars, 5 µm for D-064, 10 µm for others.

## Movies

1. Rotating view of 3D reconstruction of DHPC-018 cell, to go with Figure 1E. Cell stained with anti-Tom20 (green, mitochondria), anti-tubulin (blue), TRITC-phalloidin (red), and DAPI (white). Scale bar, 10 µm.
2. Rotating view of 3D reconstruction of a CSP, to go with Figure 1I. Cell stained with anti-CK19 (green), anti-tubulin (magenta), TRITC-phalloidin (red), and DAPI (blue). Scale bar, 10 µm.
3. Pull-away TMT assembly in which multiple TMTs are created initially, then condense to one TMT, to go with Figure 3D. Scale bar, 20 µm.
4. Pull-away TMT assembly in which one TMT is assembled initially, to go with Supplementary Figure 3A. Scale bar, 50 µm.
5. Search-and-capture TMT assembly to go with Figure 3E. Scale bar, 20 µm.
6. TMT dynamics, including ruffling, filopodium formation, and particle movement, to go with Figure 3F. Scale bar, 10 µm.
7. CSP dynamics, including ruffling at CSP tip, to go with Figure 3G. Scale bar, 10 µm.
8. CSP ruffling, filopodia, and retraction, to go with Figure 3H. Scale bar, 10 µm.
9. Rotating view of 3D reconstruction of section from a DHPC-018 flank tumor in which 10% of the injected cells express GFP (green), to go with Figure 6B. Cells stained with DAPI (blue).
10. Rotating view of 3D reconstruction of KPC 4662-derived tumor, to go with Figure 6D. Cells stained with DAPI (blue) and anti-CK19 (green).
11. Rotating view of 3D reconstruction of primary human PDAC tumor D-038, to go with Figure 6F. Cells stained with DAPI (blue) and anti-CK19 (green).

## Materials and Methods

### Culture cells

The DHPC-018 cell line was developed from a peritoneal metastasis that had been surgically removed from a 59-year old male (D11129-008) at the Dartmouth-Hitchcock Medical Center. Prior to surgery, the patient had received three cycles of gemcitabine/docetaxel/capecitabine treatment. Post-surgical testing revealed the tumor to contain the K-Ras G12R mutation. The tumor was minced, then cultured in DMEM +10% FBS +1% penicillin/streptomycin (all from Hyclone Inc) on tissue culture plastic. Cells were passaged every 3 days by 0.05% trypsinization with 0.53 mM EDTA (Corning, 25-052-Cl). After 10 passages, cells were frozen. Upon thawing, cells were maintained in DMEM (Corning, 10-013-CV) + 10% FBS (Sigma-Aldrich, F4135) for a maximum of 35 passages. KPC4662 cells (Kras^G12D/+^, Trp53^R172H/+^, Pdx1-Cre) pancreatic acinar cells were obtained from Robert Vonderheide (Winograd *et al.*, 2015) and cultured in DMEM/F12 (Corning, 15-090-CV) +10% FBS +1x Glutamax (Gibco, 35050-061) +1x Penicillin-Streptomycin (Sigma, P4333) to a maximum of 35 passages. All cultures were tested every 3 months for mycoplasma contamination using the LookOut kit (Sigma, MP0035).

To generate a stable GFP-expressing line, DHPC-018 cells were infected at passage 20 with lentivirus expressing FUGW-eGFP (Addgene, #14883), then sorted on a FACS Aria II cell sorter for GFP fluorescence. Following cell sorting, the GFP-DHPC-018 cells were cultured as the DHPC-018 wildtype cells.

### Antibodies and dyes

The following primary antibodies and dyes were used: anti-alpha tubulin (Sigma, T9026, used at 1:10000), anti-Tom20 (Santa Cruz, 11415, used at 1:300), anti-cytokeratin 19 (Abcam, 52625, used at 1:1000 for IF and 1:100 for IHC), phalloidin-tetramethylrhodamine B isothiocyanate (Sigma, P1951, used at 1:500), DAPI (Sigma, D9542, used at 1:500), and wheat germ agglutinin-647 (Molecular probes, W32466, used at 0.1 mg/mL). For immunofluorescence, the following secondary antibodies were used: anti-rabbit FITC (Invitrogen, F2765, used at 1/500), anti-mouse AF405 (Invitrogen, A31553, used at 1/500), anti-mouse AF647 (Life technologies, A21236, used at 1/500). Tissue sections were mounted in VectaShield containing added DAPI (Vector laboratories, H-1200).

### Cell plating and inhibitor treatments

Unless stated otherwise, for all cell experiments cells were seeded at 3×10^5 cells in 1.5 mL of media onto 10 µg/mL fibronectin (Sigma-Aldrich, F1141) coated MatTek dishes (MatTek Corporation, P35G-1.5-14-C) and left to incubate 16-24 hrs before treatments, fixation, or live imaging. For inhibitor treatments, cells were treated with the following: DMSO (Life technologies, D12345) at 0.5%, latrunculin A (EMD Millipore, 428021) at a 5 µM dilution from a 1 mM stock in DMSO, nocodazole (Sigma-Aldrich, M1404) at a 50 µM dilution from a 33 mM stock in DMSO, CK666 (Sigma-Aldrich, SML0006) at a 100 µM or 200 µM dilution from a 20 mM stock in DMSO, and Cytochalasin D (Sigma-Aldrich, C8273) at a 1 µM dilution from a 2 mM stock in DMSO.

### Immunofluorescence

Cells were fixed on MatTek dishes with 1% glutaraldehyde (Electron Microscopy Sciences, 16019) in BRB80 (80 mM PIPES, pH 6.9; 1 mM MgCl_2_; 1 mM EGTA) at 23°C, quenched with 3 × 10-min treatments of 1 mg/mL sodium borohydride in PBS, permeabilized with 0.25% triton in PBS for 15 min, blocked with 10% normal calf serum (HyClone, SH30118.03) in PBS with 0.02% sodium azide, and stained with antibodies and dyes in 250 µL primary or 300 µL secondary antibodies/dye incubations for 60 min each. When used, WGA-647 was administered for 5 minutes after quenching and before permeabilization. Dishes were stored in PBS at 4°C and imaged in PBS. Imaging was conducted on an LSM 880 equipped with a 63x 1.4 NA Apochromat oil objective using the Airyscan detector (Carl Zeiss Microscopy). For DAPI and AF405 secondary antibodies the 405 nm diode laser and broad pass (BP) 420-480 + BP 495-550 filter were used. For FITC and AF488 secondary antibodies the 488 nm Ar-ion laser and BP 420-480 + BP 495-550 filter were used. For rhodamine phalloidin the 561 nm diode-pumped solid state laser and BP 420-480 + 495-620 filter were used. For WGA-647 the 633 nm HeNe laser and BP 570-620 + long pass 645 filter were used. Images were acquired using Zen Black 2012 software and processed with Imaris 8.3.1 software.

### Live-cell imaging

MatTek dishes were imaged on a Nikon Ti inverted epifluorescence microscope utilizing a 20x 0.75 NA dry plan apo and 63x 1.40 NA oil plan apo objectives. DIC images were acquired with NIS Elements v5.11.02 and an Andor Zyla sCMOS FLASH 4.0 v3 camera while cells were in a TokaiHit stage-top incubator (37° C and 5% CO_2_) in DMEM + 10% FBS culture media. To maintain proper humidity over extended imaging times, a MatTek dish lid was modified by replacing a cut square region with a glass coverslip, allowing the dish to be imaged with the lid on.

### Tumor samples and immunohistochemistry

Flank tumors from the DHPC-018 cell line were generated in NOD/SCID mice following IACUC protocol #2072, allowing 6 weeks for tumor growth. For most preparations, mice were euthanized by CO_2_ asphyxiation, then tumors were excised and fixed in 10% formalin, 4% paraformaldehyde (Electron Microscopy Sciences, 15710), 1% glutaraldehyde (Electron Microscopy Sciences, 16000), or a combination of 1% glutaraldehyde and 4% paraformaldehyde in PBS. Fixed tumors were either sectioned immediately on a vibratome (50 μm sections) and stored in PBS with 4% sucrose until staining, or paraffin-embedded and sectioned later (FFPE, 20 μm sections). For one mouse, fixation was performed by cardiac puncture using 4% PFA (Acros Organics, AC416780250) and 4% sucrose (Fisher, BP220-1) in PBS (National Diagnostics, CL-253) fixative (following IACUC protocol #0002073(m15)), then the excised tumor was sectioned immediately (50 μm sections). For FFPE samples, deparaffinization was conducted in xylene (EMD Millipore, XX0055), followed by hydration in graded ethanol/water, then the tumor underwent antigen retrieval in boiling TRIS-EDTA buffer (10 mM Tris, 1 mM EDTA, pH 9) for 15 min before blocking in 4% Triton-X100 with 10% normal calf serum (Hyclone, SH30118.03) in PBS, staining with anti-CK19 (where stated) and mounting in DAPI-containing VectaShield. Non-embedded samples were subject to the same procedures, omitting the deparaffinization, hydration, and antigen retrieval steps. For assessment of tumors directly in mouse pancreas, tumors were created by orthotopic injection of KPC4662 cells. Briefly, 5×10^5-1×10^6 cultured KPC4662 cells (Kras^G12D/+^, Trp53^R172H/+^, Pdx1-Cre) pancreatic acinar cells (Winograd *et al.*, 2015) are injected into the pancreatic parenchyma (below the capsule) of C57BL/6 mice. At four-weeks post-injection, the mouse is euthanized by CO_2_ asphyxiation, the pancreas is excised and subject to FFPE preparation, sectioning and staining as described above. Primary human tumors were obtained under IACUC protocol 2177 (see Table 1) as FFPE blocks. 20 μm tumor sectioning and staining were prepared as for mouse FFPE sections above.

Tumors were imaged on an LSM880 Airyscan confocal microscope. Tile-scans were performed using either a 10x 0.3 NA or 20x 0.8 NA dry objective (depending on the size of the tissue section), acquiring a series of 3 µm Z steps and 16-50+ individual fields which were stitched together using Zen Black image processing prior to Airyscan processing. Higher resolution images of tumor TMTs were performed using a 63x 0.75 NA oil objective, acquiring a series of 0.2 µm Z steps at regions with cells staining positive for CK19.

### Quantification of TMT and CSP dimensions and frequency

TMT/CSP frequency and length were determined from 20x z-stacks acquired on the Dragonfly microscope (10-25 slices at 1 μm). Fields were acquired randomly throughout the dish. For frequency, the number of cells, TMTs and CSPs were counted in the entire imaging field (Andor Zyla sCMOS FLASH 4.0 v3 camera, ??? × ??? μm area). Length was measured for all TMTs and CSPs throughout the field. TMTs were counted if they were at least 2 slices above the basal surface, and CSPs were counted if they originated at least 2 slices above the basal surface. TMT and CSP width and height above basal surface were measured from 63x z-stacks acquired on the Airyscan microscope (15-40 slices at 0.4 μm). Width was determined using WGA staining, at the z-slice containing the widest TMT section. Line-scans were taken across the TMT, and the exterior position at half height of each side of the TMT was used to mark the position of the plasma membrane.

### Quantification of TMT assembly frequency

For Figure 3B, 3.5×10^5 cells in 1.5 mL of media were seeded onto 10 µg/mL fibronectin coated MatTek dishes and left in the incubator for 4 hours. After incubation, five fields of DIC images were acquired on a Nikon Eclipse Ti microscope, 20x 0.75 NA objective, with a time interval of 30 min for 24 hrs. During the first 3 hrs, every new TMT assembly was counted and categorized as either a pull-away or search-and-capture event. For Figure 5C, cells were seeded at 3×10^5 cells in 1.5 mL of media onto 10 µg/mL fibronectin coated MatTek dishes and left to incubate 16-24 hrs before treatment with DMSO or 200 µM CK666. Dishes were imaged with the same microscope and objective as above, with the following changes: three fields were acquired per experiment with 15 min intervals for 3 hrs.

## Acknowledgments

We would like to thank the PDX Mouse Service at the Norris Cotton Cancer Center, Robert Vonderheide (University of Pennsylvania) for supplying the KPC4662 cell line, Tim Bruceos for forming connections, and Shivaprasad Sathyanarayana for help with perfusion fixation and vibratome sectioning. This work was supported by NIH R35 GM122545, a seed grant from the Hirshberg Foundation for Pancreatic Cancer Research, and a Prouty Pilot Project grant to HNH; NIH R01 CA204228-01 to SDL; a Steven B. Currier Clinical Oncology Scholar Award to KDS, NCI Cancer Center Support Grant 5P30CA023108-37 to the Norris Cotton Cancer Center; and NIH P20 GM113132.

